# Genetically Encoded FerriTag as a Specific Label for Cryo-Electron Tomography

**DOI:** 10.1101/2024.09.10.612178

**Authors:** Chang Wang, Ioan Iacovache, Benoît Zuber

## Abstract

Cryo-electron tomography (cryoET) is an important imaging technique that can provide 3D datasets of organelles and proteins at nanometer and sub-nanometer resolution. Recently, combining cryoET with subtomogram averaging has pushed the resolution to 3-4 Å. However, one main challenge for cryoET is locating target proteins in live cells. Conventional methods such as fluorescent protein tagging and immunogold labeling are not entirely suitable to label small structures in live cells with molecular resolution in vitrified samples. If large proteins, which can be visually identified in cryoET, are directly linked to the target protein, the large tag may alter the target protein structure, localization and function. To address this challenge, we used the rapamycin-induced oligomer formation system, which involves two tags (FKBP and FRB) that can bind together within rapamycin. In our system, the FKBP tag is linked to target protein and the FRB tag is linked to a large protein to create a marker. We chose ferritin as the marker protein because it is a large complex (10–12 nm) and can bind iron to create strong contrast in cryoET. After adding rapamycin to the cell medium, the iron-loaded ferritin accurately indicates the location of the target protein. Recently, in-situ cryoET with subtomogram averaging has been rapidly developing. However, it is still challenging to locate target proteins in live cells, and this method provides a much-needed solution.

## Introduction

CryoET is a powerful imaging method that enables the visualization of cellular structures in their native state at nanometer resolution (Gan & Jensen, 2012). By combining cryoET with subtomogram averaging, researchers have recently pushed the achievable resolution to 3-4 Å, allowing for unprecedented insights into cellular architecture and macromolecular complexes (Xing et al., 2023 and von Kügelgen et al., 2024). CryoET is attracting significant interest. However, a persistent challenge in cryoET is the precise localization of proteins of interest within the complex cellular milieu.

Conventional methods for protein labeling that are in use in fluorescence microscopy or in electron microscopy (EM) of resin-embedded samples (conventional EM) are not entirely suitable for cryoET applications. While correlative light and electron microscopy (CLEM) approaches exist, the localization precision is often insufficient to unambiguously assign proteins to specific structures observed in the tomograms with nanometer precision (Guaita et al., 2022). Furthermore, immunogold labeling, a technique used in EM of resin-embedded samples, poses significant challenges for intracellular structures in cryoET. This is because cryoET requires all labeling steps to be performed on native cells (i.e., not fixed, not permeabilized) prior to vitrification. Gold-coupled antibodies or other large probes cannot easily be introduced into intact cells without compromising cellular integrity, thus limiting their applicability for intracellular targets in cryoET studies. Additionally, diaminobenzidine (DAB) oxidation, another widely used method in conventional EM, is not applicable in cryoET because the visible product relies on the aggregation of osmium, which cannot be used in cryoET sample preparation (Martell et al., 2017). These constraints highlight the need for alternative labeling strategies that are compatible with the unique requirements of cryoET while maintaining cellular and structural integrity.

Recent advancements have shown promise in the use of gold nanoparticles for cryoET labeling. Groysbeck et al. demonstrated that 2 nm gold particles functionalized with nuclear localization signals can be introduced into cultured cells via electroporation and subsequently imported into the nucleus (Groysbeck et al., 2023). This approach overcomes some of the limitations of traditional gold labeling methods. However, this technology still requires significant development to achieve specific protein labeling. Moreover, the small size of these 2 nm beads, while advantageous for cellular uptake, presents challenges for detection in cryoET datasets. The limited contrast of such small particles makes them difficult to distinguish from cellular structures in the complex intracellular environment visualized by cryoET.

Ferritin, a large protein complex (∼450 kDa) forming a hollow cage of 12 nm in diameter, has emerged as a promising candidate for protein labeling in EM (Jutz et al., 2015). It can store up to 4500 iron atoms in its 8-nm diameter core, resulting in electron-dense particles that are easily identifiable in EM. Ferritin offers several advantages over alternative labeling methods. It possesses a highly symmetrical structure with remarkable chemical and thermal stability (Khoshnejad et al., 2018). The ferritin cage can be reconstituted through controlled reassembly (Pead et al., 1995), making it suitable for various experimental conditions. Furthermore, a broad range of metals can be loaded and mineralized inside the ferritin cavity, providing flexibility in contrast generation (Jutz et al., 2015). Both the interior and outer surface of the ferritin cage can easily be modified through the addition of peptides or protein tags, enabling versatile functionalization. Finally, ferritins are highly biocompatible and less immunogenic than other protein cages, reducing potential artifacts in cellular systems (Kim et al., 2016).

The potential of ferritin as a labeling marker in cryoET was first demonstrated by Wang et al., (Wang et al., 2011). In their pioneering work, they overexpressed FtnA, a bacterial ferritin, linked to GFP along with target protein, such as membrane protein, chemosensory machinery and septum proteins in *Escherichia coli*. The location was first identified by the fluorescence of GFP-FtnA, followed by visualization of the ferritin tag in cryoET. This method showed low cell toxicity and provided good visibility of the tag. However, a serious drawback of this approach is that fusing ferritin directly with the protein of interest can lead to strong artifacts because ferritin, and consequently the protein of interest, oligomerize. This oligomerization can significantly alter the native behavior and localization of the target protein, potentially compromising the biological relevance of the observations.

To address this limitation and provide a more versatile labeling approach, Clarke and Royle developed a more sophisticated ferritin labeling system called FerriTag (Clarke & Royle, 2018). This innovative approach leverages the rapamycin-induced dimerization system, combining the FKBP (FK506 binding protein) and FRB (FKBP-rapamycin binding) domains to overcome the challenges associated with direct ferritin fusion (Inobe & Nukina, 2016; Spencer et al., 1993). In this system, FRB is linked to ferritin, while FKBP is fused to the target protein along with a fluorescent protein such as GFP. FRB and FKBP can rapidly dimerize in the presence of rapamycin, a bacterial antifungal molecule. Upon addition of rapamycin, the ferritin-FRB complex binds to the FKBP-tagged target protein, accurately indicating its location. This approach allows for temporal control of labeling and combines the advantages of fluorescence microscopy for initial localization with the high-resolution capabilities of electron microscopy for precise structural analysis. Clarke and Royle have applied their system in combination with EM of resin-embedded samples to localize mitochondrial outer membrane proteins as well as clathrin. Their results suggest that the system may be applicable to cryoET, since all labeling steps are performed prior fixation.

Recently, Fung et al. developed a similar approach using the rapamycin-induced dimerization system coupled with genetically encoded multimeric particles (GEMs) for protein labeling in cryoET. Their system employs icosahedral particles derived from encapsulins, ranging from 25 to 42 nm in size, which are coupled to GFP-tagged proteins of interest upon addition of rapamycin. This method has been successfully applied to label various cellular targets, including mitochondrial, nuclear, and endoplasmic reticulum proteins, demonstrating its versatility for subcellular protein localization in cryoET studies (Fung et al., 2023).

In our study, we chose to adapt the FerriTag system due to its smaller label size and enhanced visibility when loaded with iron. The compact nature of this probe allows it to access more tightly packed cellular regions, enabling labeling in areas where larger tags might be excluded. We have further improved upon this system by constructing a simple plasmid design, which simplifies the labeling procedure. While a recent preprint by Sun et al. (Sun et al., 2024) has shown the use of iron-free FerriTag for cryoET in membrane patches obtained by cell unroofing, our approach goes further. Importantly, we have demonstrated the visibility of the FerriTag in vitreous sections of high-pressure frozen cells studied by cryo-electron microscopy of vitreous sections (CEMOVIS), as well as in cryoET of focused ion beam (FIB)-milled, plunge-frozen cells (Studer et al., 2014). Our approach has been successfully applied to label several outer mitochondrial membrane proteins and KRas, a protein that associates with the inner leaflet of the plasma membrane through lipid anchors. These results underscore the versatility and effectiveness of our labeling strategy across different cellular compartments and membrane-associated proteins. Our work paves the way for the identification and structural characterization of proteins in their native three-dimensional cellular context with molecular resolution, offering unprecedented insights into the complex organization and interactions within living cells.

## Materials and methods

### Cell culture

Cells were cultured in a humidified incubator at 37 °C with an atmosphere containing 5% CO_2_. HEK293T cells were grown in Dulbecco’s modified Eagle’s medium (Gibco, Thermo Fisher Scientific) supplemented with 10% FCS (BioConcept) and penicillin-streptomycin (Gibco). 2 x 10^6^ cells were frozen in 500 µl cell-freezing media (Bambanker, BB01). After thawing at 37 °C in a water bath for 1 min, cells were diluted 10-fold with cell culture media and centrifuged for 5 min at 1000 x g. Subsequently, the cells were seeded into a 10 cm dish with 15 ml cell culture media. After 24 hours, the cells were passaged once.

For staining with Mito Tracker™ Deep Red (Thermo Fisher Scientific, M22426), 1 mM stock solution in DMSO was prepared, and a final working concentration of 500 nM was used. The solution was added to cell culture medium for 15 minutes at 37 °C. Following incubation, the cells were fixed in 4% paraformaldehyde and observed by confocal microscopy (excitation wavelength range 644-665 nm).

### Plasmids

All plasmids used in this study are listed in Table 1. Plasmids of FRB-mCherry-FTH1, FKBP-GFP-Myc-MAO and untagged FTL were a kind gift of Prof. Stephen Royle. FKBP-mCherry-BclXLc33 was a gift from Ivan Yudushkin (Addgene plasmid # 72905; http://n2t.net/addgene:72905; RRID:Addgene_72905) (Ebner et al., 2017), FKBP-mCherry-KRas4Bc30 was a gift from Ivan Yudushkin (Addgene plasmid # 72900; http://n2t.net/addgene:72900; RRID:Addgene_72900) (Ebner et al., 2017). Tom20-mCherry-FKBP was a gift from Takanari Inoue (Addgene plasmid # 162436; http://n2t.net/addgene:162436; RRID:Addgene_162436) (Wu et al., 2020). FRB-FTL-IRES-FTH and FTL-IRES-FRB-mCherry-FTH were cloned into plasmid pLVX-M-puro, which was a gift from Boyi Gan (Addgene plasmid # 125839; http://n2t.net/addgene:125839; RRID:Addgene_125839) (Zhang et al., 2018).

### Transfection

When cells reached 80% confluency, plasmids were transfected using Lipofectamine™ 3000 (L3000008, Thermo Fisher Scientific). For 3.5 cm dish, 4 μL Lipofectamine™ 3000 was diluted in 125 μL of Opti-MEM™ in one tube. In another tube, 750ng of FKBP-GFP-MAO, 600ng of FTL and 150ng of FRB-mCherry-FTH1 were diluted into 125 μL Opti-MEM™, followed by the addition of 5 μL P3000™ reagent to the diluted plasmid mixture. Then, the two tubes were mixed and then incubated for 15 mins at room temperature. After incubation, the mixture was added dropwise to the cells. The treated cells were ready for use after 2 days post-transfection. For transfection of FRB-FTL-IRES-FTH and FKBP-GFP-MAO, 750ng of each plasmid was used. For transfection of PLVX-FTL-IRES-FRB-mCherry-FTH-PGK-FKBP-GFP-MAO, 1.5μg of plasmid was used.

### Immunofluorescence microscopy

Cells were fixed in 4% paraformaldehyde for 10 min at room temperature on a shaker. Following fixation, the cells were washed 3 times with phosphate-buffered saline (PBS) and permeabilized with 0.5% Triton X-100 in PBS, gently shaking for 5 mins. Coverslips were then incubated with primary antibody (Ferritin light chain, Proteintech, 10727-1-AP) at a 1:200 dilution in 10% bovine serum albumin (BSA)-PBS at 4 °C overnight. Afterward, the coverslips were washed three times with PBS and incubated with secondary antibody (Goat-anti-rabbit, Biorbyt, orb705814 which has an excitation peak at 651 nm and an emission peak at 670 nm, diluted 1:200 in 10% BSA-PBS) for 45 mins at room temperature. After three PBS washes, the slides were treated with an antifade mounting medium containing DAPI (Invitrogen™S36968). The fluorescence was observed using confocal microscopy (Zeiss LSM 880 with Airyscan, 63x 1.4 NA oil objective).

### Sample preparation for conventional transmission electron microscopy

Cells were transfected with plasmid using Lipofectamine™ 3000 and treated with 1 mM FeSO_4_·7H2O (Sigma, F8633) supplemented medium for 16 hours (Clarke & Royle, 2018). After a 15-minute rapamycin treatment, cells were fixed in 3% glutaraldehyde in 0.05 M phosphate buffer at pH 7.4 for 1 h. Following fixation, cells were washed 3 times in 0.05 M phosphate buffer and then post-fixed in 1% osmium tetroxide (Agar) for 1 h. After three washes, they were dehydrated through a series of ethanol solutions (70%, 80%, 96%, 100%), with each step lasting 15 min. In the end, the cells were embedded using 50% epoxy resin (TAAB) in ethanol 2 hours and then 100% epoxy resin (TAAB) overnight at room temperature. The cells were then polymerized in 100% epoxy resin at 60 °C for 3 days.

### Sample preparation for high pressure freezing and CEMOVIS

Cells were collected and mixed in a volume ratio of 1:1 with 40% dextran in PBS (70 kDa; Sigma–Aldrich, product number: 31390). This mixture was then inserted into copper tubes as previously described (Studer et al., 2014), and sample vitrification was achieved by high pressure freezing in an EM PACT2 (Leica Microsystems). After vitrification, a Leica EM UC6 ultramicrotome with an EM FC6 cryochamber was operated at a temperature of −150 [. The extremity of the tube was trimmed away with a trimming diamond knife (45° Diatome, Nidau, Switzerland), using a feed of 200 nm and a speed of 100 mm/s until the entire sample surface showed even blackness (indicating no holes on the surface and no ice crystals within the sample). Cryosection ribbons were generated with another diamond knife (35° diamond knife, Diatome) with a feed of 50 nm and a speed of 1 mm/s. An ionizer was used in discharge mode during sectioning and in charge mode for attaching the ribbon onto the EM grid (Lacey Carbon Film 200 Mesh Cu, Agar Scientific) using micromanipulators (Studer et al., 2014).

### Sample preparation for plunge freezing

HEK293T cells were transfected with the respective plasmids. After 24 hours, cells were seeded onto holey carbon gold EM grids (200 mesh, R2/2, Quantifoil) coated with 10 mg/ml fibronectin. Simultaneously, 1 mM iron was added to cell medium. After 16 hours, rapamycin (final concentration 200 nM) was added to the medium. Following a 15-minutes incubation, HEPES at pH 7.2 (final concentration: 10 mM) was added to cell medium, and the grids were immediately plunge-frozen. Grids were manually blotted on the back-side for 20 seconds with Whatman filter paper No.1, and then vitrified in liquid ethane. Frozen grids were screened for ice quality and cells of interest were identified using cryo-FM in a Leica EM cryo-CLEM system (Leica Microsystems) under a humidity-controlled atmosphere.

### Cryo-FIB milling and cryoET

Lamellae were generated through cryo-FIB milling using an Aquilos 2 Cryo FIB (Thermo Fisher Scientific), following a procedure similar to that published previously (Franken et al., 2022; Ganeva et al., 2023). The grids were subjected to gas injection system coating for 90 s at a working distance of 10 mm, resulting in a platinum layer of approximately 1 μm thickness. Scanning electron microscopy (SEM) was used to locate cells of interest at 5 kV voltage and 13 pA current and for imaging to verify milling at 2 kV voltage and 13 pA current. For rough milling, the voltage was maintained at 30 kV in all steps. The milling steps were as follows: 1) 1 nA, 35° stage tilt, milling a lamella of 20 μm thickness; 2) 0.5 nA, 25° tilt, milling to 12 μm thickness; 3) 0.3 nA, 17° tilt, reducing the thickness to 3 μm; and 4) 0.1 nA, 17° tilt, further thinning to 1 μm. For the fine milling steps, the voltage was reduced to 16 kV and the current to 23 pA. Initially, the stage was tilted to 18° and the lamella was milled from the front side, then the stage was tilted to 16°, and the lamella was milled from the back side. After these two steps, the lamella thickness was approximately 0.5 μm. Finally, the stage was tilted back to 17°, and the current was adjusted to 11pA. The lamella was milled from the top and the bottom to achieve a target thickness approximately 0.2 μm.

CryoET data of FIB-milled cells was acquired on a Krios G4 cryo-transmission electron microscope at 300 kV voltage, with a Falcon 4i direct electron detector and a Selectris energy filter (Thermo Fischer Scientific). Tomo program (Thermo Fisher Scientific) was used for acquisition. During acquisition the stage was set to tilt from −60° to +60° (initially tilting the stage to the angle of the lamellae) and each picture was acquired per angle. The dose of each tilt series image was adjusted to 1∼1.2 e^-^/Å^2^ and the target dose rate at the detector was kept around 4 e^-^/px/s. For data analysis, IMOD was used (Kremer et al., 1996).

### Protein purification and analysis

Expi293F cells were obtained from Thermo Fisher Scientific (Cat. No. A14527). Cells were cultured in 250 mL flasks containing 60 mL of Expi293 Expression Medium (Thermo Fisher Scientific, Cat. No. A1435101). Cells were maintained at a density of 2 × 10^6^ cells/mL, incubated at 37°C with 8% CO2, and shaken at 125 rpm. For transfection, one of two plasmids was used: either His-FTH1 or His-FRB-FTL-IRES-FTH1. A total of 76 µg of the selected plasmid DNA was diluted in 1.5 mL Opti-MEM. Separately, 240 µL of polyethylenimine (PEI, 1 mg/mL; Polysciences Inc., Cat. No. 24765-1) was diluted in 1.5 mL Opti-MEM. The diluted DNA and PEI solutions were mixed and incubated for 20 minutes at room temperature. The resulting transfection mixture was then added dropwise to the cell culture. Cells were harvested 48 hours post-transfection for protein purification.

Cell pellets were resuspended in lysis buffer containing PBS, 20 mM imidazole, and 2 mM β-mercaptoethanol (β-ME), supplemented with EDTA-free protease inhibitor cocktail and benzonase (Sigma-Aldrich). Cell lysis was performed by sonication using 4 cycles (10 seconds on at 35% power, 60 seconds off) at 0°C. The cell lysate was centrifuged at 25,000 rpm for 1 hour using a 45 Ti rotor. The supernatant was loaded onto a 5 mL Ni-NTA column (GE Healthcare) connected to an ÄKTA Prime plus for purification.

Protein samples were vitrified for cryo-EM analysis as described above. Initial screening of samples was performed using a FEI Tecnai F20 cryo-electron microscope.

Single-particle data were collected using a Krios G4 cryo-electron microscope as described above. Data acquisition was automated using EPU software (Thermo Fisher Scientific). We picked particles and performed 2D classification using CryoSPARC (Punjani et al., 2017).

## Results

We sought to adapt the FerriTag system, previously developed for conventional EM labeling, for cryoET applications (Clarke & Royle, 2018). This system employs a rapamycin-induced ferritin-target oligomer formation mechanism, potentially allowing for specific labeling in non-chemically fixed, vitrified samples. The key advantage is that until rapamycin addition, the target protein function is minimally affected, being fused only to the small FKBP tag (∼12 kDa).

To validate the iron-loading capacity of overexpressed ferritin, we first purified His-tagged FTH1 from cells cultured with and without iron supplementation. CryoEM analysis of these purified samples revealed clear differences in iron content within the ferritin cages based on iron availability in the culture medium (Supplementary Figure 1A). Quantitative analysis showed that approximately 45% of ferritin cages contained visible iron when cells were cultured with 1 mM iron, compared to only about 10% in the absence of iron supplementation (Supplementary Figure 1B). 2D classification of single-particle cryoEM data of ferritin purified from cells that had not been treated with iron further illustrated the varying cation content within ferritin particles, with loaded ferritin showing distinctive contrast in their core regions compared to empty particles (Supplementary Figure 1C). These results confirm that overexpressed ferritin effectively incorporate iron under physiological conditions, providing a strong basis for their use as an electron-dense tag in subsequent labeling experiments.

### Labeling the mitochondria for CEMOVIS

To assess the labeling capabilities of our FerriTag system, we chose monoamine oxidase (MAO) as our target protein. MAO is located in the mitochondrial outer membrane and an enzyme that catalyzes the oxidative deamination of monoamines (Cohen & Kesler, 1999). This aids in validating the labeling method, as mitochondria are easily identifiable structures in cryoEM images. We linked FKBP to MAO and FRB to monomeric ferritin. To validate the mitochondrial localization of FKBP-GFP-MAO we used MitoTracker™ Deep Red, a reliable mitochondrial fluorescent marker. HEK293T cells were transfected with FKBP-GFP-MAO as described in the methods. After 24 hours, cells were passaged to new dish. After cells reached 30% confluency, they were treated with MitoTracker™ Deep Red, followed by rapamycin addition. From confocal microscopy results, the fixed cells revealed strong colocalization of FKBP-GFP-MAO and MitoTracker™ Deep Red labeled mitochondria (Figure 1A), confirming the successful targeting of our construct to mitochondria.

**Figure 1.**
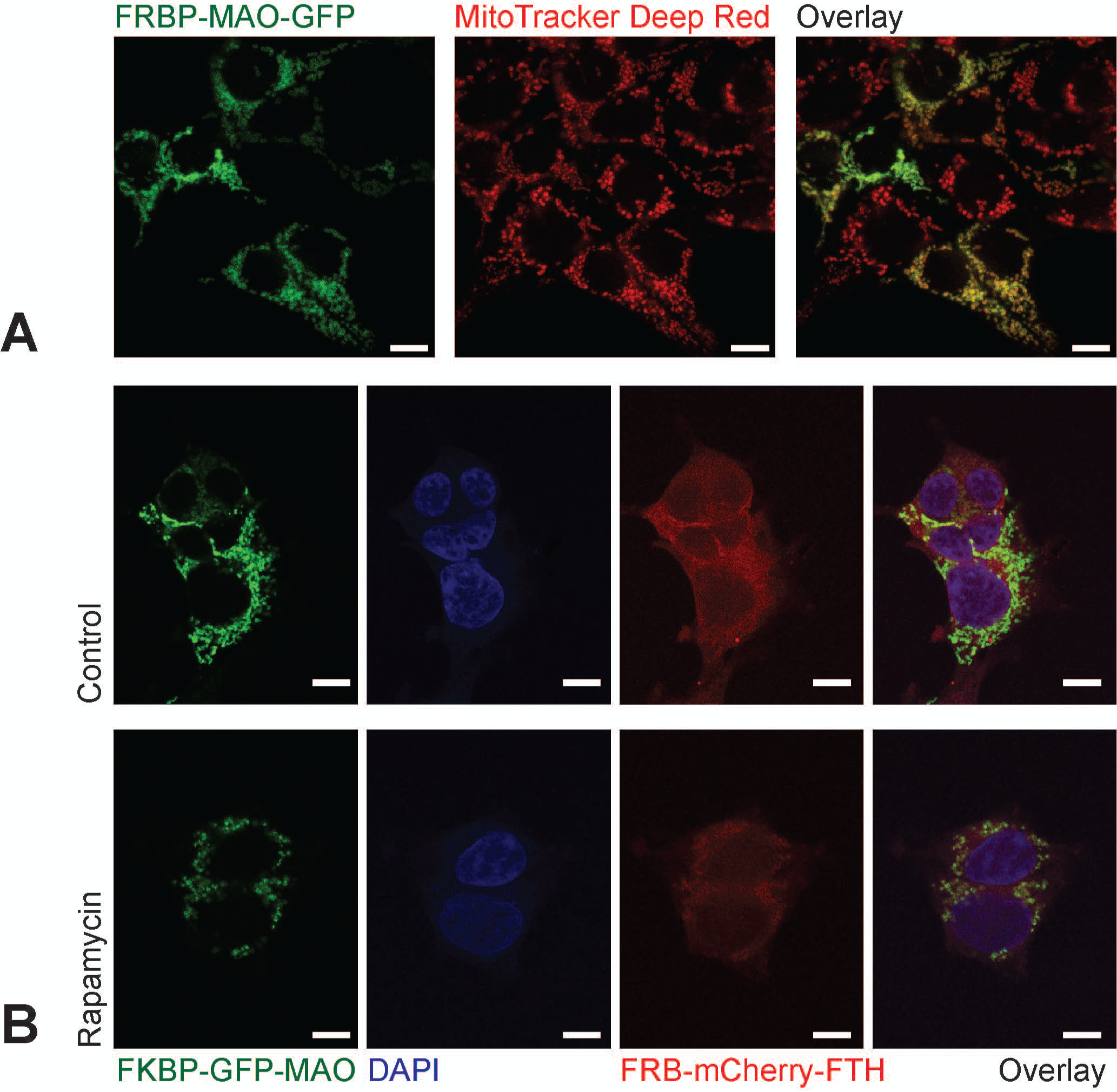
Rapamycin-induced ferritin labeling of MAO using the three-plasmid system. **A**. Cells transfected with FKBP-MAO-GFP were treated with MitoTracker Deep Red. The FKBP-MAO-GFP is expressed and correctly targeted to the mitochondria as shown by the colocalization with MitoTracker. **B**. Cells were co-transfected with three plasmids, FKBP-MAO-GFP, FRB-mCherry-FTH and untagged FTL. In the control sample FRB-mCherry-FTH is distributed throughout the cytoplasm while FKBP-MAO-GFP is seen as before on the mitochondria. DAPI was used to stain the nucleus. Upon addition of rapamycin (200nM for 15 minutes) FKBP-MAO-GFP and FRB-mCherry-FTH colocalized on the mitochondria. Scale bars: 10 µm.

We next evaluated the rapamycin-induced association with the target protein. HEK293T cells were co-transfected with FKBP-GFP-MAO, FRB-mCherry-FTH (ferritin heavy chain) and untagged FTL (ferritin light chain). From confocal microscopy, in the presence of rapamycin FKBP-GFP-MAO localized to mitochondria, while mCherry-FTH was diffusely distributed throughout the cytoplasm (Figure 1B). Upon rapamycin treatment, we observed a significant colocalization of FRB-mCherry-FTH and FKBP-GFP-MAO (Figure 1B). This colocalization demonstrates successful rapamycin-induced dimerization of the FKBP and FRB tags, effectively recruiting ferritin to MAO at the mitochondrial outer membrane.

To visualize the ferritin labeling at higher resolution, we examined cells expressing the same constructs using conventional EM. In cells treated with rapamycin prior fixation, we observed many electron-dense particles specifically located at the surface of mitochondria (Figure 2A). In contrast, control cells not treated with rapamycin showed a diffuse distribution of these particles throughout the cytoplasm, with no specific association with mitochondria (Figure 2A). This difference in ferritin localization further confirms the efficacy of the FerriTag system.

**Figure 2.**
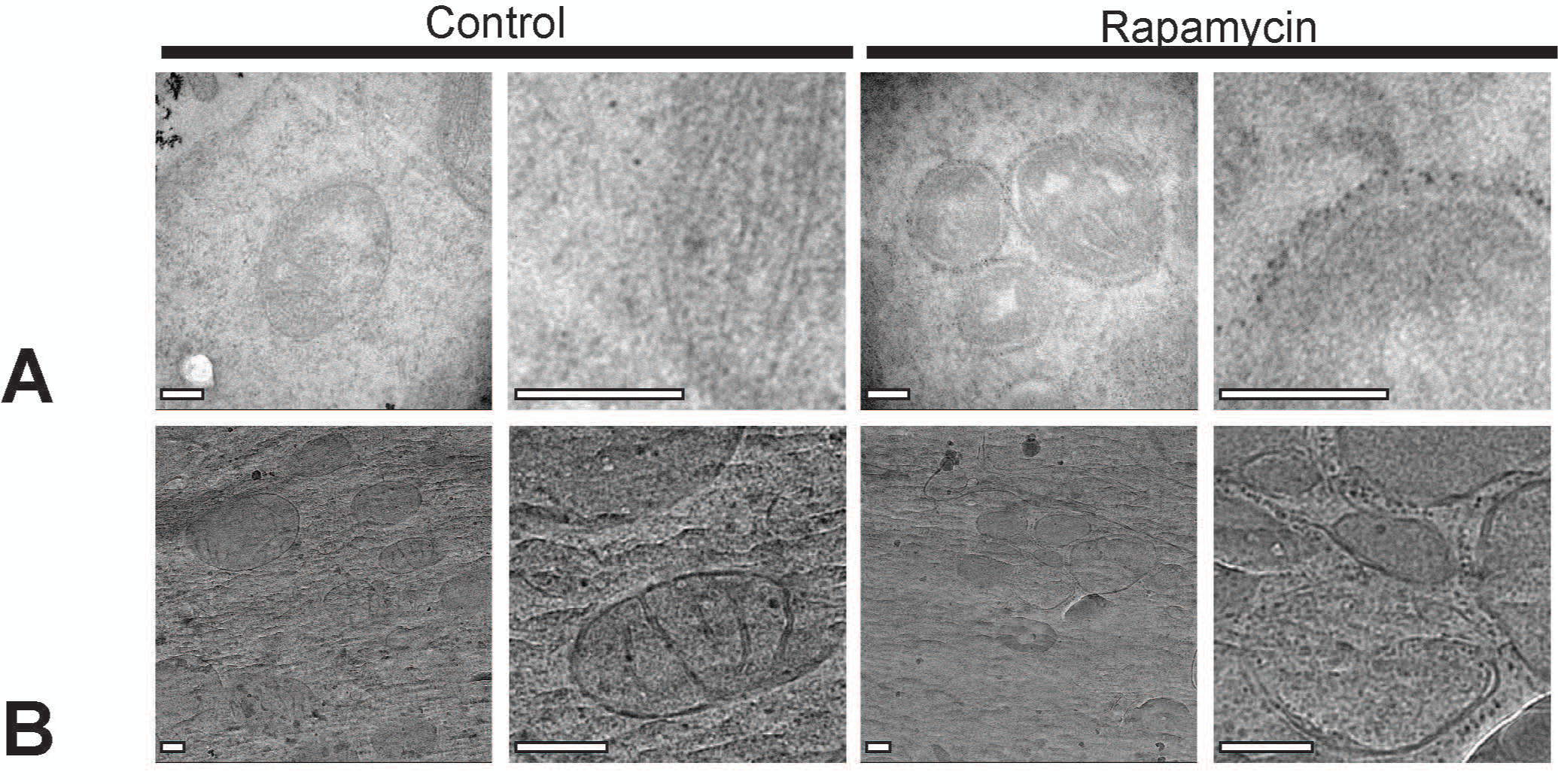
TEM visualization of rapamycin-induced labeling of Mao by Ferritin in TEM using the three-plasmid system. **A**. Cells were co-transfected with three plasmids as in Figure 1B (FKBP-MAO-GFP, FRB-mCherry-FTH, and untagged FTL) and 1mM iron (II) sulfate was added to the cell media. The cells were fixed, stained and sectioned for TEM with or without prior rapamycin addition. In the absence of rapamycin, small electron-dense particles are seen in the cytoplasm while the mitochondrial membrane is unlabeled. Upon rapamycin addition the mitochondrial membrane is covered with electron-dense particles, confirming the efficacy of the FerriTag system. **B**. Cells grown as in **A** were vitrified by high pressure freezing followed by cryo-sectioning by CEMOVIS. The thin sections were imaged by cryo-fluorescence to identify the areas of interest. As before in the absence of rapamycin the mitochondrial membrane shows no ferritin labeling while upon rapamycin treatment the mitochondria is surrounded by electron-dense particles. Scale bars: 200 nm.

Our primary objective was to observe iron-loaded ferritin labeling in vitrified samples, preserving cellular structures in their native state. To achieve this, cells were vitrified by high-pressure freezing and processed into ribbons of ultrathin vitreous sections ribbons by cryo-ultramicrotomy (CEMOVIS). We employed a correlative approach, first using cryo-fluorescence microscopy to locate regions with FKPB-GFP-MAO and FRB-mCherry-FTH signals in the ribbons. These regions were then examined by cryoEM. We observed 7-10 nm electron-dense particles surrounding the mitochondrial outer membrane, consistent with iron-loaded ferritin (Figure 2B). when rapamycin was omitted, we did not observe this specific decoration of mitochondrial membranes with electron-dense particles.

The results from both conventional EM and CEMOVIS demonstrate that the FerriTag system provides specific and readily identifiable labeling in native cellular context, confirming its potential as a versatile marker for cryoEM and cryoET.

### Optimization of FerriTag plasmid design for efficient single-vector expression

To enhance the versatility of the FerriTag system, we sought to simplify the labeling procedure, making it convenient for targeting diverse proteins. To achieve efficient expression of multiple components from a single vector, we employed internal ribosome entry site (IRES) for polycistronic expression and phosphoglycerate kinase (PGK) promoter for robust transcription. We designed two plasmids: FTL-IRES-FRB-mCherry-FTH and FTL-IRES-FRB-mCherry-FTH-PKG-FKBP-GFP-MAO. Since genes positioned upstream of the IRES sequence typically show higher expression levels (Kazadi et al., 2008), we placed FTL before the IRES and FRB-mCherry-FTH after it. This design ensures higher expression of untagged FTL compared to FRB-mCherry-FTH, promoting the assembly of ferritin oligomers with an excess of untagged ferritin subunits, as recommended by (Clarke & Royle, 2018). The PGK promoter was used to drive robust expression of the target protein.

We transfected cells with the FTL-IRES-FRB-mCherry-FTH plasmid and the FKBP-GFP-MAO at a 1:1 ratio. The cells were imaged using both confocal microscope and conventional EM. Consistent with our previous observations, confocal imaging showed colocalization of FRB-mCherry-FTH and FKBP-GPF-MAO in the presence of rapamycin (Figure 3A). It confirmed that the IRES-mediated expression of FRB-mCherry-FTH was successful and that FKBP-GFP-MAO bound to FRB-mCherry-FTH upon rapamycin addition. Conventional EM further corroborated these findings, showing iron-loaded ferritin particles surrounding mitochondria after rapamycin treatment (Figure 3A), as expected.

**Figure 3.**
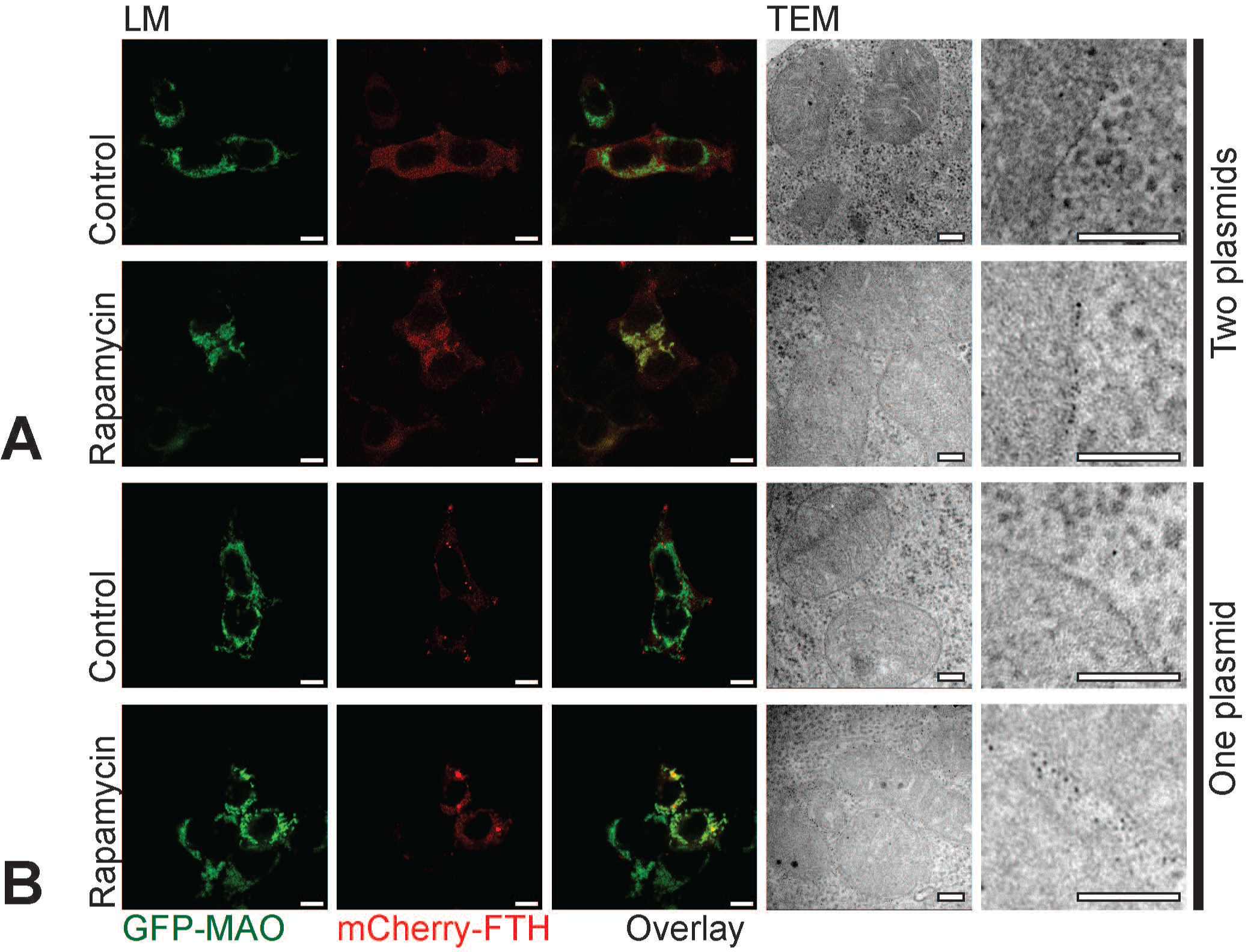
Optimizing the labeling of MAO by Ferritin. **A**. Two plasmids (FTL-IRES-FRB-mCherry-FTH and FKBP-GFP-MAO) were transfected into cells at a 1:1 ratio and the cells were imaged as before by both confocal microscopy and TEM. As in the three-plasmid system we could observe colocalization of GFP-MAO and mCherry-FTH by fluorescence and could confirm labeling by TEM. **B**. A single plasmid (FTL-IRES-FRB-mCherry-FTH-PKG-FKBP-GFP-MAO) was transfected to assess the expression of all components from one plasmid. Fluorescence showed clear colocalization between GFP-MAO and mCherry-FTH upon rapamycin addition and the results were corroborated by TEM imaging where the mitochondria is seen surrounded by electron dense particles upon rapamycin addition. Scale bars: 10 µm for confocal microscopy, 200 nm for EM.

Next, we assessed the expression of all components incorporated into a single plasmid (FTL-IRES-FRB-mCherry-FTH-PKG-FKBP-GFP-MAO). Following transfection and rapamycin treatment, we observed colocalization of FKBP-GFP-MAO and FRB-mCherry-FTH (Figure 3B), indicating the single plasmid successfully expressed FTL, FRB-mCherry-FTH, and FKBP-GFP-MAO in cells. These results were comparable to those obtained with the initial three-plasmid system (Figure 2) and the subsequent two-plasmid approach (Figure 3A). Conventional EM revealed iron-loaded ferritin surrounding the mitochondria (Figure 3B), consistent with our previous observations. However, when we examined these samples using cryoEM, we identified only a few properly labeled sites. This unexpected result could be attributed to several factors. One possibility is that the expression level of FRB-mChery-FTH was insufficient to achieve robust labeling of mitochondria upon rapamycin addition. Alternatively, the ferritin may not be properly assembled and/or may not be properly loaded with enough iron, reducing its visibility in cryoEM.

### Optimizing FerriTag for enhanced cryoET labeling efficiency

To address the lower ferritin signal observed in cryoET, we redesigned our plasmids to increase protein expression. We modified the construct by linking FRB linker to FTL instead of FTH and positioned the FRB-FTL fusion upstream of the IRES. This new design led to increased production of iron-loaded ferritin, generating stronger labeling signal. Additionally, to eliminate any potential interference with ferritin oligomerization, we removed the mCherry tag from FTH.

We first validated our new constructs by examining the colocalization of FKBP-MAO-GFP and FTL, detected by immunofluorescence, upon rapamycin treatment (Figure 4A). To assess suitability of our optimized FerriTag for protein labeling and localization in cryoET, we prepared samples by high pressure freezing and cryo-ultramicroscopy. We then used correlative cryo-fluorescence microscopy to identify positive samples for subsequent cryoEM imaging. With these newly designed plasmids, we clearly observed the ferritin tags surrounding the mitochondria after rapamycin treatment (Figure 4B). In addition to CEMOVIS, we explored the use of cryoFIB, a recently established technique for producing thin lamellae suitable for cryoET. Unlike CEMOVIS, which produces ribbons containing dozen of sections, each covering roughly 25,000 µm^2^, cryoFIB generates much smaller areas. A typical cryoFIB lamella is only about 50 µm^2^ in area, with about a dozen lamellae produced per sample. This dramatic reduction in area of electron-transparent regions makes precise localization of the target proteins crucial for successful cryoFIB milling. Consequently, correlative light and electron microscopy is essential to guide the milling process. We therefore tested our new FerriTag construct in a cryoFIB workflow that incorporates cryo-fluorescence microscopy to identify regions of interest before milling and cryoET imaging. Implementing this workflow, we milled 200 nm thick lamellae at locations identified by cryo-fluorescence microscopy in the vitrified samples. The lamellae were then re-inspected by cryo-fluorescence microscopy to verify that the labeled proteins were not milled away and to precisely localize them. When examined by cryoEM and cryoET, these samples showed improved image quality compared to CEMOVIS preparations, which suffered from cutting artifacts. Notably, we clearly observed ferritin dots surrounding mitochondria, demonstrating the effectiveness of our optimized FerriTag system in cryoFIB-prepared samples (Figure 4B, Supplementary movie 1 and 2).

**Figure 4.**
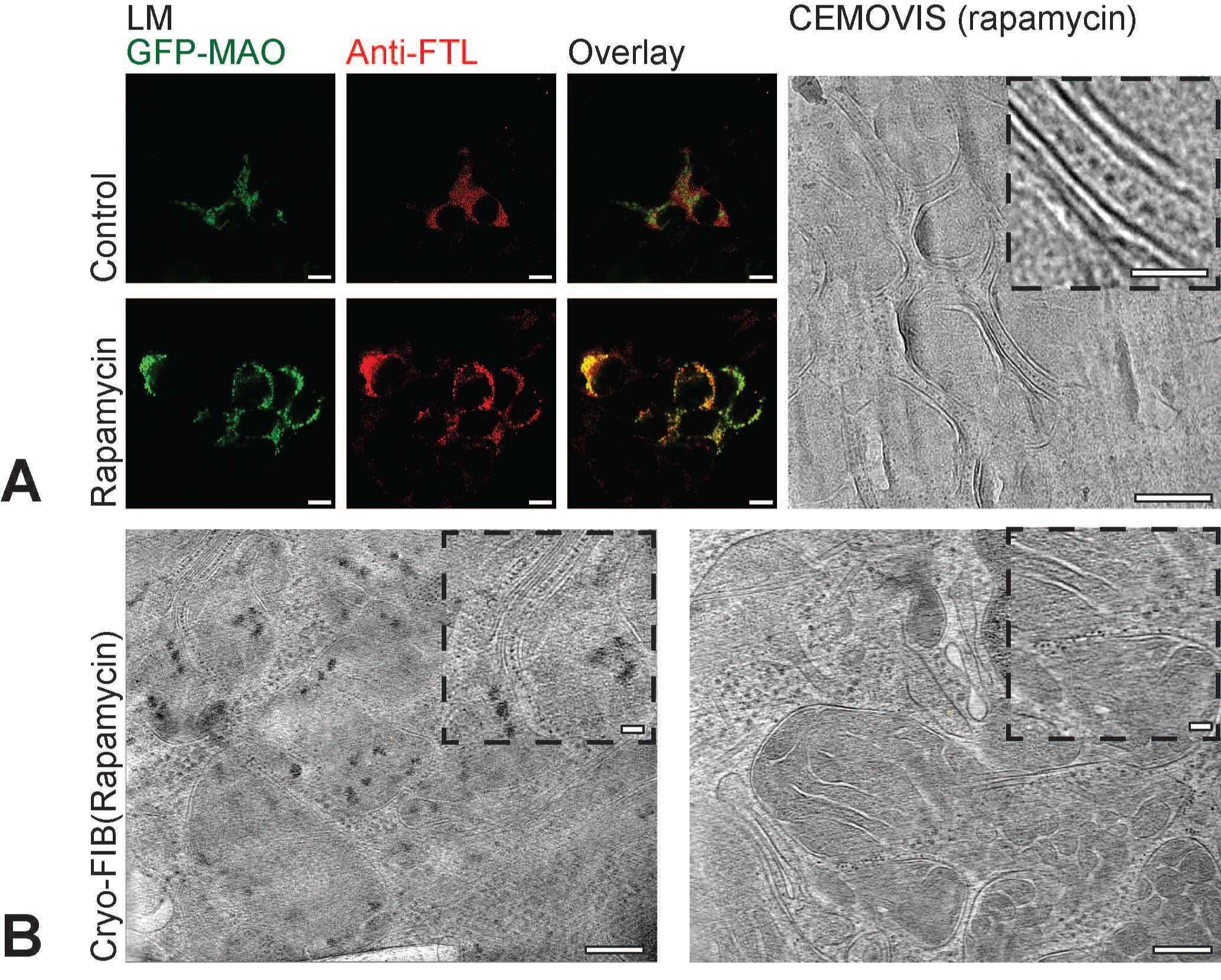
Optimizing the labeling of MAO by Ferritin for cryo-EM and cryo-ET. **A**. Two plasmids (FKBP-GFP-MAO and FRB-FTL-IRES-FTH) were transfected into cells. Fluorescence confirmed correct expression of GFP-MAO and FTL and showed colocalization of FTL and MAO upon rapamycin addition. Following vitrification by high pression freezing and CEMOVIS the ferritin labeling was shown to correctly localize to the mitochondrial membrane upon rapamycin addition. **B**. The sample was vitrified by manual blotting and plunging in liquid ethane and cryo-FIB was used to prepare a 200 nm lamella. Ferritin is seen labeling the mitochondrial membrane as before upon rapamycin addition. Scale bars: 10 µm for confocal microscopy, 200 nm for EM, 50 nm for magnified insets.

### Versatility of FerriTag: Labeling diverse proteins

To demonstrate the versatility of our optimized FerriTag system, we targeted three additional proteins: TOM20, Bcl-xL and K-Ras. TOM20 is a mitochondrial outer membrane protein that recognizes and facilitates the translocation of precursor proteins from the cytoplasm to mitochondria (Likić et al., 2005). Bcl-xL is also an anti-apoptotic mitochondrial outer membrane protein (Finucane et al., 1999). K-Ras is a plasma membrane-associated GTPase, involved in signal transduction (Kranenburg, 2005).

Similar to our MAO labeling experiments, we observed that FKBP-mCherry-KRAS and FRB-FTL colocalized after rapamycin treatment, as evidenced by confocal microscopy. When examined by conventional EM, we clearly observed iron-loaded ferritin around the plasma membrane. We then demonstrated the effectiveness of FerriTag in cryogenic conditions using CEMOVIS. In rapamycin-treated cells, we detected iron-loaded ferritin at the plasma membrane, accurately reflecting the known location of KRAS (Figure 5A). Following rapamycin treatment, we demonstrated that both TOM20 and Bcl-xL could be labeled with ferritin, as observed by confocal microscopy. We then examined the samples using conventional EM, which revelaled iron-loaded ferritin around the mitochondria for both proteins as expected. CEMOVIS analysis yielded similar results, clearly showing iron-loaded ferritin surrounding the mitochondria for both TOM20 and Bcl-xL (Figure 5B, 5C).

**Figure 5.**
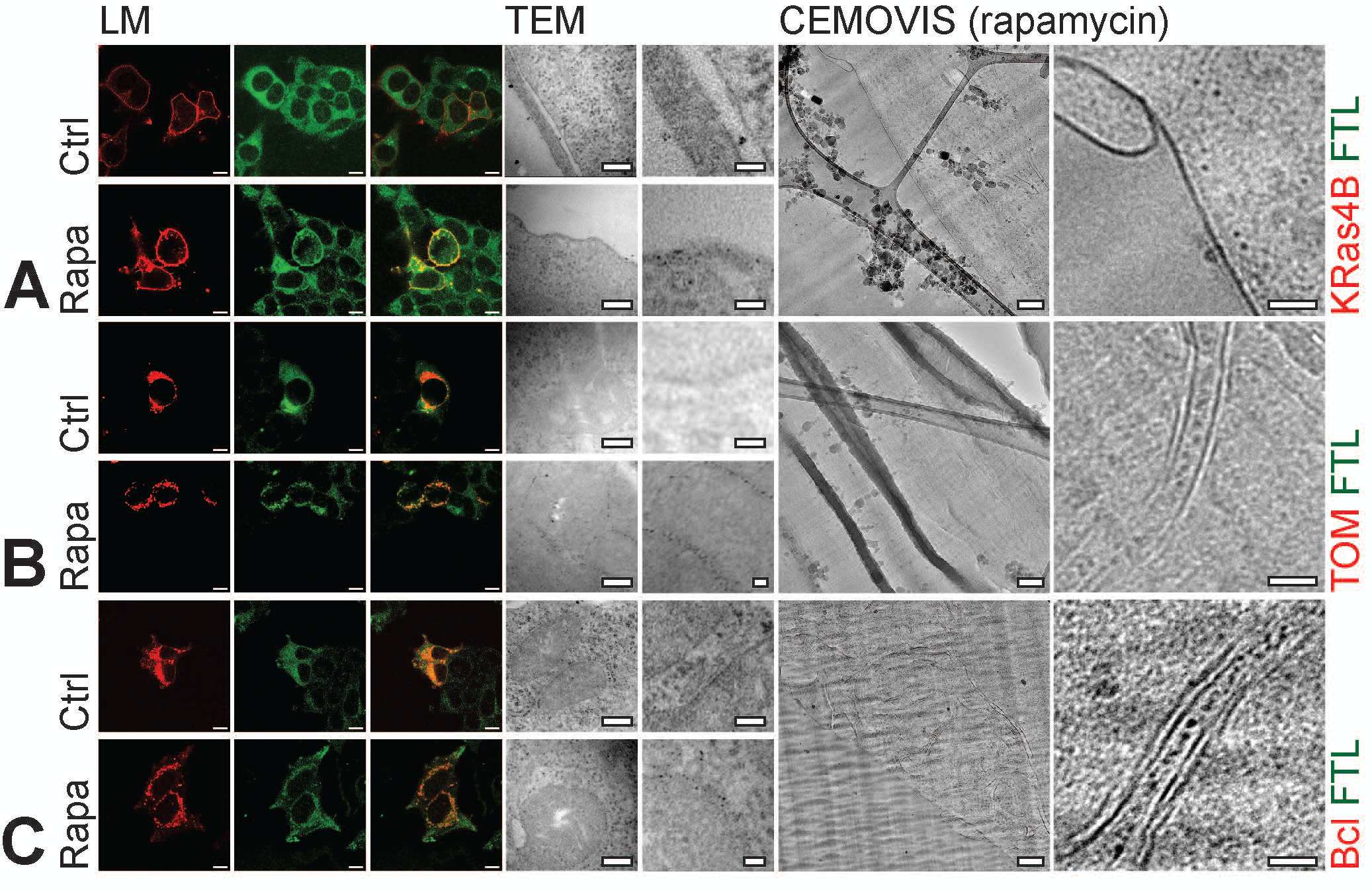
Confirming the utility of the FerriTag system for labeling in cryo-EM and cryo-ET. **A.** KRas4B (red), a plasma membrane protein, was labeled with the FerriTag system. Confocal microscopy and TEM confirmed colocalization of ferritin (green) with the plasma membrane upon rapamycin addition. Cryo-EM on CEMOVIS sections confirmed ferritin labeling of the plasma membrane upon rapamycin addition. **B**. TOM (red) and **C**. Bcl (red) were labeled with the Ferritag system and the successful labeling was confirmed by confocal microscopy, TEM as well as cryo-EM on CEMOVIS upon rapamycin addition. Scale bars: 10 µm for LM, 200 nm for EM, 50 nm for magnified rightmost images

## Discussion

In this study, we sought to develop and optimize a versatile protein labeling method for correlative fluorescence and cryoEM. Our primary objective was to overcome the limitations of existing labeling techniques in cryoET, where traditional methods such as immunogold labeling or fluorescent protein tags often fall short. The key innovation of our approach lies in the adaptation and optimization of the FerriTag system (Clarke & Royle, 2018; Park et al., 2021), originally developed for conventional electron microscopy, for use in cryogenic conditions.

Our work has resulted in several significant outcomes. Firstly, we developed a simple and efficiency FerriTag system, which streamlines the transfection process and improves the efficiency of protein labeling. This system combines the ferritin components (FTL and FTH) with FRB module in a single construct. We found that this single-plasmid approach, as well as a dual-plasmid system separating the ferritin components, can be effectively used for conventional EM applications.

Second, we demonstrated the efficacy of our optimized FerriTag system in cryoET. The iron-loaded ferritin tags provided strong contrast and specificity, allowing for precise localization of target proteins within the cellular ultrastructure under cryogenic conditions. This represents a significant advance in protein labeling for cryoET, where traditional labeling methods often fall short.

Lastly, we demonstrated the versatility of our labeling approach by successfully targeting proteins in different cellular compartments. We achieved specific labeling of mitochondrial outer membrane proteins (MAO, TOM20, and Bcl-xL) as well as a plasma membrane-associated protein (K-Ras). This demonstrates the broad applicability of our method across various cellular structures and protein types, underlining its potential as a valuable tool for diverse structural biology studies.

The FerriTag system offers several advantages for protein labeling in cryoET. Unlike immunogold labeling, it allows for in situ labeling in living cells prior to vitrification, preserving cellular context. The ferritin tag provides a clear, electron-dense signal that is easily identifiable in tomograms, overcoming the limitations of fluorescent proteins in cryoET.

The rapamycin-inducible system allows for temporal control of labeling, minimizing artifacts associated with constitutive expression of large tags. The small size of the FKBP tag (∼12 kDa) reduces functional interference prior to labeling. In cryoET, the 12 nm ferritin particles, with their 8 nm dense core, strike a balance between visibility and accessibility, being large enough to identify yet small enough to access most cellular compartments.

Compared to the GEMs described by Fung et al. (Fung et al., 2023), both systems achieve similar localization precision. Our FerriTag system offers advantages in size (12 nm vs. 25-42 nm) and iron-ferritin were visible in cryoEM pictures. Our approach shares similarities with the preprint study of Sun et al. (Sun et al., 2024b) in using FerriTag but differs in key aspects of sample preparation. Both systems achieve similar localization precision. Our approach shares similarities with the preprint study of Sun et al. in using FerriTag but differs in key aspects of sample preparation. Both studies employ FerriTag in living cells before any sample preparation for cryoET.

However, while Sun et al. subsequently unroof their cells to create membrane sheets before vitrification, we maintain the integrity of the whole cell through the vitrification process. This allows us to preserve the complete cellular architecture, including the cytoplasmic context of membrane-associated proteins. Interestingly, our study and Sun et al work differ in their approach to FerriTag preparation. While we use iron-loaded FerriTag, Sun et al. demonstrate the effectiveness of iron-free FerriTag in cryoET. Since they work with membrane patches, their samples are very thin and iron-free FerriTag are readily visible. Our use of iron-loaded FerriTag provides strong contrast and easy identification in the cellular environment. Nonetheless, Sun et al. work suggests that iron-free FerriTag might also be detected in intact cells, potentially making this labeling approach applicable to cell types that are sensitive to or do not tolerate iron supplementation. Detection of iron-free FerriTag in the complex environment of intact cells could be further enhanced by employing AI-based particle recognition methods, similar to those used by Fung et al. for the GEM particles, thereby improving the efficiency and accuracy of protein localization in cryoET.

The FerriTag system for cryoET has significant implications for cellular biology. While cryoET allows us to visualize cellular structures at molecular resolution, identifying specific proteins within these structures remains challenging. FerriTag addresses this by enabling precise protein identification, thus linking observed structures to their functions. As cryoET techniques continue to improve, the FerriTag approach will play a crucial role in developing comprehensive, functionally annotated maps of cellular architecture. Ultimately, this method provides a valuable tool for investigating the molecular basis of cellular functions, offering new insights into fundamental biological processes and potentially contributing to our understanding of various health and disease states.

## Acknowledgements

This work was supported by the Swiss National Science Foundation (grant numbers 31003A_179520, CRSII_222809, and 32NE30_185536 to BZ). We gratefully acknowledge the Dubochet Center for Imaging - Bern (DCI-Bern) and the Microscopy Imaging Center (MIC) of the University of Bern for their technical support and access to advanced imaging facilities. We extend our sincere thanks to Wanda Kukulski and her group for their advice regarding FIB-milling. We also thank Stephen Royle, Ivan Yudushkin, Takanari Inoue, and Boyi Gan for sharing plasmids.

## Author contributions

BZ conceived and supervised the project. CW prepared samples, acquired and processed the data. CW and II analyzed the data. All authors wrote the manuscript.

## Ethics declarations

The authors declare no competing financial interests.

## Supplementary Information

**Supplementary Figure 1**

**Supplementary Movie 1**

**CryoET of FerriTag-labeled mitochondria in FIB-milled lamella.** This movie shows a tomogram of a 200 nm-thick cryo-FIB lamella from cells expressing FRB-FTL-IRES-FTH1 and FKBP-MAO. The electron-dense ferritin particles are clearly visible surrounding the mitochondrial membranes.

**Supplementary Movie 2**

**CryoET of FerriTag-labeled mitochondria in FIB-milled lamella.** The sample was prepared as in Movie 1. Several patches of ferritin are visible at the surface of mitochondria.

**Supplementary Movie 3**

**CryoET of mitochondria with reduced FerriTag expression.** The sample was prepared as in Movie 1, except for the FRB-FTL-IRES-FTH1 plasmid concentration, which was halved. Ferritin particles are particularly abundant in the lower part image. Arrows points to some ferritin-containing areas

