## Supplementary Figure 1 for "Genetically Encoded FerriTag as a Specific Label for Cryo-Electron Tomography"

**Supplementary Information**

**Supplementary Figure**

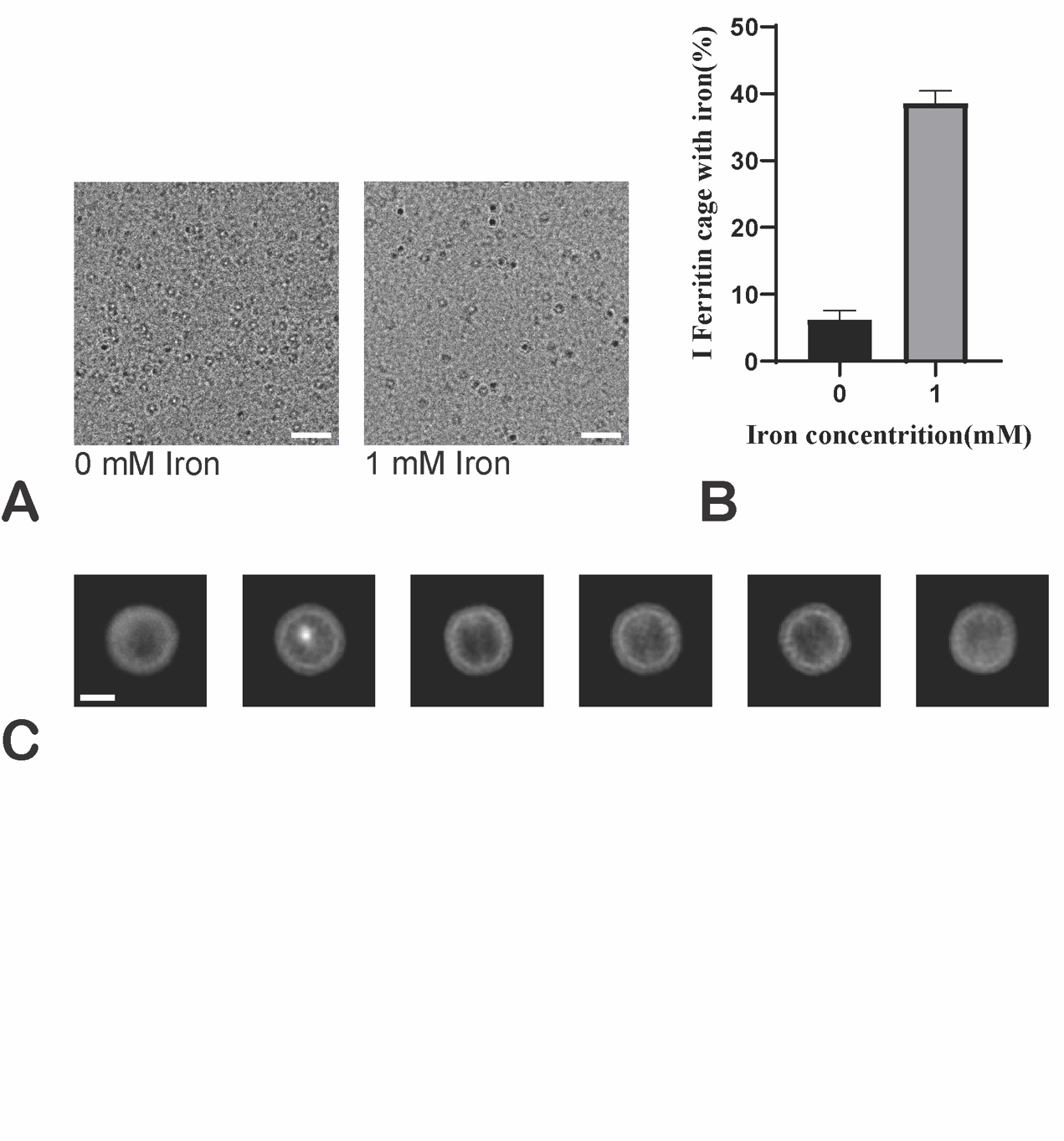

**Supplementary Figure 1**

The his-FTH1 was overexpressed in HEK293T cells for 24 hours and then incubated with 1mM FeSO4·7H2O in cell medium. After 24 hours of iron incubation, the cells was lysed to purify the ferritin protein. CryoEM images shown, approxiately 40% of ferritin cage were capable of binding iron (Supplementary figure A and B). Additionally, we purified the his-FRB-FTL-IRES-FTH1 and 2D classification from the CryoSPARC analysis revealed iron at the core of ferritin cage (Supplementary figure C).
